# Annotation-free prediction of microbial dioxygen utilization

**DOI:** 10.1101/2024.01.16.575888

**Authors:** Avi I. Flamholz, Joshua E. Goldford, Elin M. Larsson, Adrian Jinich, Woodward W. Fischer, Dianne K. Newman

**Affiliations:** Division of Biology and Biological Engineering, California Institute of Technology, Pasadena, CA 91125; Division of Geological & Planetary Sciences, California Institute of Technology, Pasadena, CA 91125; Skaggs School of Pharmacy and Pharmaceutical Sciences, University of California at San Diego, La Jolla, CA 92093; Department of Chemistry and Biochemistry, University of California at San Diego, La Jolla, CA 92093

**Keywords:** Biogeochemistry, oxygen physiology, machine learning, large language models

## Abstract

Aerobes require dioxygen (O_2_) to grow; anaerobes do not. But nearly all microbes — aerobes, anaerobes, and facultative organisms alike — express enzymes whose substrates include O_2_, if only for detoxification. This presents a challenge when trying to assess which organisms are aerobic from genomic data alone. This challenge can be overcome by noting that O_2_ utilization has wide-ranging effects on microbes: aerobes typically have larger genomes, encode more O_2_-utilizing enzymes, and tend to use different amino acids in their proteins. Here we show that these effects permit high-quality prediction of O_2_ utilization from genome sequences, with several models displaying >70% balanced accuracy on a ternary classification task wherein blind guessing is only 33.3% accurate. Since genome annotation is compute-intensive and relies on many assumptions, we asked if annotation-free methods also perform well. We discovered that simple and efficient models based entirely on genome sequence content — e.g. triplets of amino acids — perform about as well as intensive annotation-based algorithms, enabling the rapid processing of global-scale sequence data to predict aerobic physiology. To demonstrate the utility of efficient physiological predictions we estimated the prevalence of aerobes and anaerobes along a well-studied O_2_ gradient in the Black Sea, finding strong quantitative correspondence between local chemistry (O_2_:sulfide concentration ratio) and the composition of microbial communities. We therefore suggest that statistical methods like ours can be used to estimate, or “sense,” pivotal features of the environment from DNA sequencing data.

**Importance:** We now have access to sequence data from a wide variety of natural environments. These data document a bewildering diversity of microbes, many known only from their genomes. Physiology — an organism’s capacity to engage metabolically with its environment — may provide a more useful lens than taxonomy for understanding microbial communities. As an example of this broader principle, we developed algorithms that accurately predict microbial dioxygen utilization directly from genome sequences without first annotating genes, e.g. by considering only the amino acids in protein sequences. Annotation-free algorithms enabled rapid characterization of natural samples, demonstrating a quantitative correspondence between sequences and local O_2_ levels. These results suggest that DNA sequencing can be repurposed as a multi-pronged chemical sensor, estimating concentrations of O_2_ and other key facets of complex natural settings.

## Introduction

Dioxygen (O_2_) is a hugely consequential molecule for the biosphere. Aerobic respiration yields a tremendous amount of energy and is the most common bioenergetic mode in cells across Earth’s surface environments. Yet O_2_ is also highly reactive, presenting challenges to organisms that encounter it (1). Aerobes require O_2_ to grow; anaerobes do not. But nearly all microbes — aerobes, anaerobes and facultative organisms alike — express enzymes whose substrates include O_2_, if only for detoxification. This presents a challenge when trying to assess which organisms are aerobic from genomic data alone (2, 3).

Aerobes, however, are different from anaerobes, and these differences — though subtle — are legible in their genomes. Aerobes tend to have larger genomes (4) with more O_2_ utilizing enzymes (2, 5) and usually belong to specific phylogenetic groups (4). Conversely, anaerobes make use of diverse fermentation pathways to conserve energy in low O_2_ settings (3, 5). These differences have been used to predict O_2_ utilization from the genome with reasonable accuracy (2, 6, 7).

Classification of microbial O_2_ utilization typically relies on intensive preprocessing where, for example, O_2_ utilizing enzymes are identified by sequence homology (2) or a full metabolic network is reconstructed (6). This processing is costly and limited by our very incomplete knowledge of the relationship between protein sequence and function. We therefore asked whether high-quality classification can be achieved without annotation, instead using DNA and protein sequences directly to build classifiers.

The input for classification is a genome and the output is an O_2_ utilization category. Here we considered the ternary classification problem — categorizing a genome as an (i) obligate aerobe, (ii) obligate anaerobe, or (iii) a facultative aerobe (6). Due to limited data, accuracy is typically evaluated by cross-validation — repeatedly reserving small subsets of training data for evaluation (2, 6, 7). We developed a more stringent approach, training classifiers on a compendium of ≈3300 genomes with known O_2_ utilization (8), and evaluating accuracy using an independent dataset (2).

## Observation

Genomes can be processed in different ways to enable classification (Fig 1A). A typical intensive pipeline begins with identification of protein coding sequences (ORF prediction) followed by homology-based annotation of gene functions (2, 7). Further processing is sometimes performed, for example, by constructing a metabolic network based on annotated genes (6). Processed data are then used to train classifiers that predict attributes like carbon source preference (9) or O_2_ utilization from genomic features. Features can include the counts of annotated gene functions — e.g., 1 benzoate dioxygenase, 2 heme oxygenases, etc., (2) — or the suite of molecules that could be produced by annotated enzymes (6, 9).

**Fig. 1:**
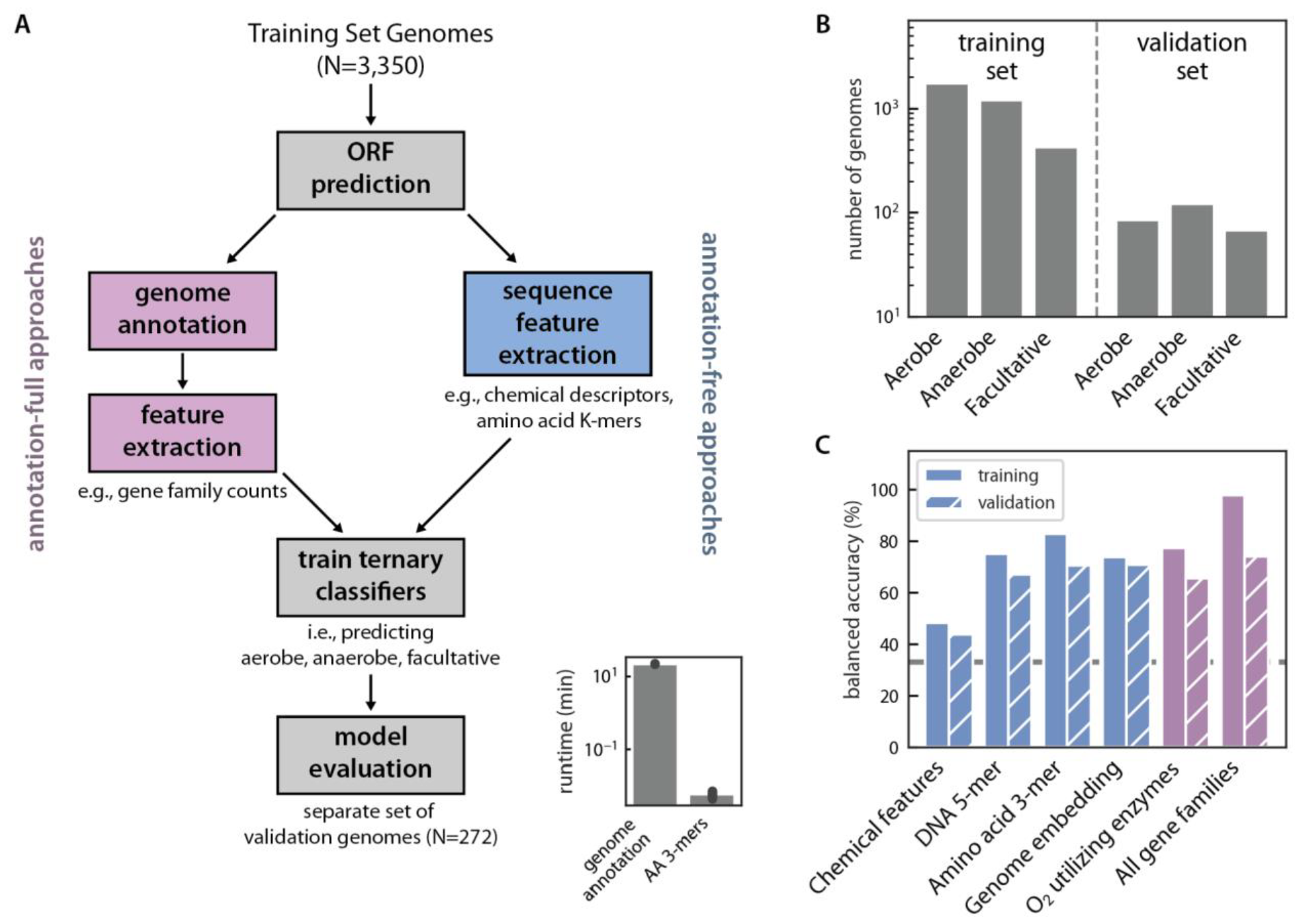
Comparing annotation-full and annotation-free approaches to predicting microbial O_2_ utilization. (A) Schematic of data processing pipelines for predicting O_2_ utilization with (left branch, purple) and without (right branch, blue) compute-intensive genome annotation. Bottom right inset: genome annotation (≈10 min/genome) is far slower than extraction of amino acid trimers (< 1 s/genome on a laptop computer). (B) The number of genomes contributing to the training (8) and independent validation (2) datasets. (C) The best-performing models were ≈70% accurate at classifying out-of-sample genomes as aerobes, anaerobes or facultative. These included an annotation-free method trained on amino acid (AA) 3-mers (70%), and a compute-intensive approach relying on whole-genome annotation to identify known protein functional groups (as defined by the KEGG database, 74% accuracy). The dashed gray line marks the expected 33% performance of picking one of three categories at random.

Avoiding annotation and working instead with nucleotide (NT) and amino acid (AA) sequences elides most pre-processing steps. Reducing processing can simplify pipelines, remove assumptions, and substantially improve runtime (Fig. 1A, inset). One simple way of representing patterns in NT and AA sequences is by counting *K*-mers — substrings of length *K* (10, 11). A more complex, and potentially valuable, annotation-free approach uses advances in machine learning to summarize (“embed”) protein sequences in vectors of fixed dimension for training classifiers (12, 13).

Comparison of model accuracies revealed that the best-performing classifier involved genome annotation (14), predicting microbes’ O_2_ utilization — obligate aerobe, obligate anaerobe or facultative — from counts of annotated gene functions (KEGG orthogroups, Fig. 1C). This model displayed 74% validation accuracy as calculated with a balanced metric that accounts for unequal representation of O_2_ utilization classes in the validation set. Yet we found that several annotation-free classifiers also exceeded 66% validation accuracy (2x the accuracy of blind guessing), including models based on counts of AA triplets (70%), NT 5-mers (67%), and protein sequence embeddings (70%).

Counting AA triplets is far more efficient than annotating genomes, which accelerated our evaluation of O_2_ utilization in environmental samples. To demonstrate the utility of rapid characterization, we analyzed ≈50,000 metagenome assembled genomes (MAGs) assembled from Earth Microbiome Project samples (15). Consistent with expectations, samples of characteristically anaerobic habitats (e.g. rumen, 603/606 MAGs classified as anaerobes) contained a much larger proportion of anaerobic MAGs than settings with higher O_2_ (e.g. freshwater lakes, 30/416, Fig. 2A).

**Fig. 2:**
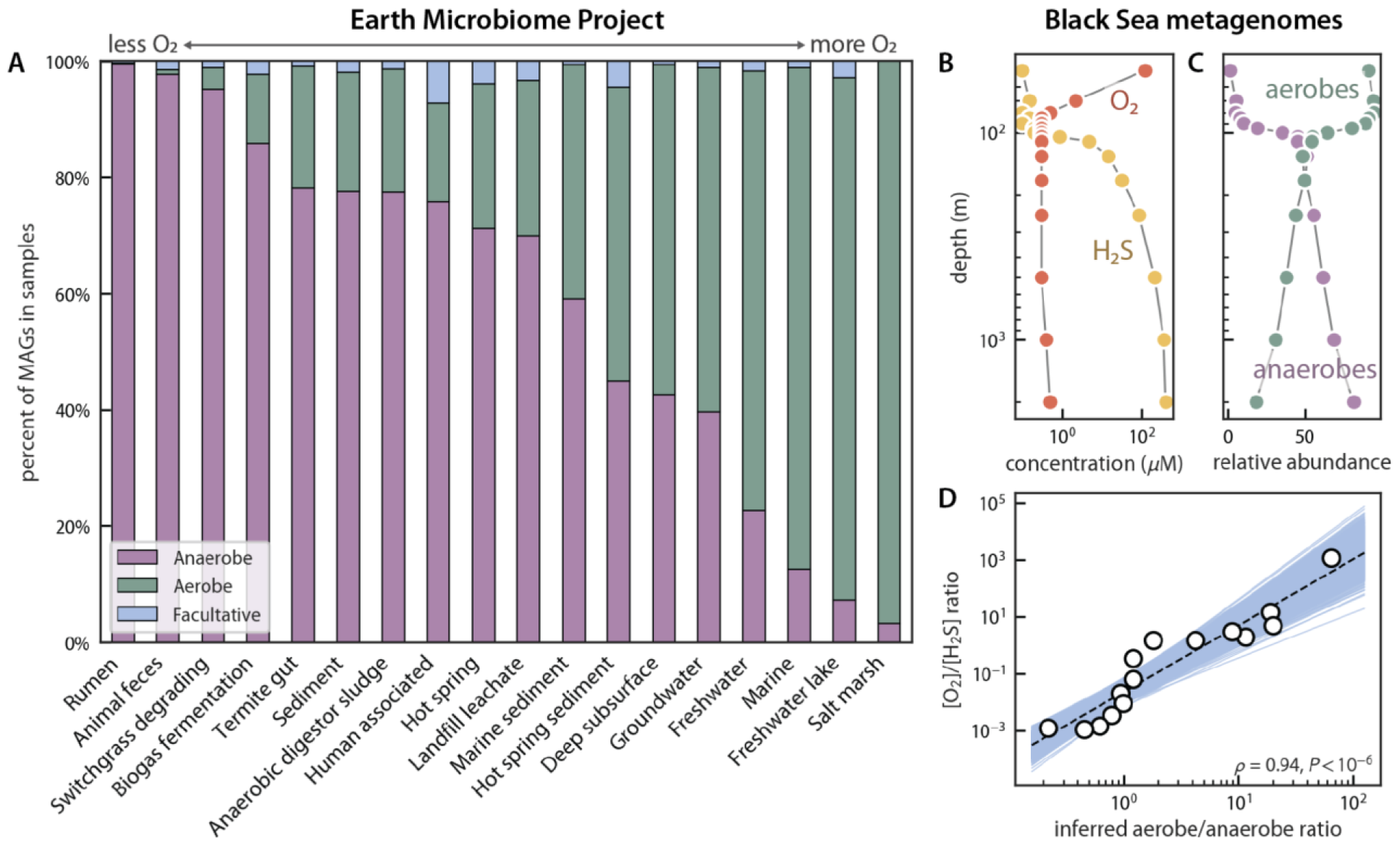
O_2_ is a major determinant of the sequence composition of natural microbial communities. (A) We applied our AA 3-mer classifier to Earth Microbiome Project samples for which sufficient numbers of metagenome assembled genomes (MAGs) were available (*n* ≥ 10). Samples from environments with characteristically low O_2_ levels (e.g. ungulate rumen, anaerobic digesters) displayed high anaerobe content, while, conversely, oxic surface environments predominantly hosted MAGs inferred to be aerobes (e.g. freshwater lakes). (B-D) The Black Sea is a well-studied stratified euxinic ecosystem with a long-lived systematic O_2_ gradient — oxygenated at the surface with a loss of O_2_ and an increase in sulfide with depth. We drew depth-dependent O_2_ and H_2_S concentrations as well as MAGs from (16). (C) We applied our annotation-free AA 3-mer model to 160 MAGs assembled in (16) to estimate the depth-dependent prevalence of aerobes and anaerobes. Consistent with O_2_ and H_2_S profiles, aerobes were most prevalent near the surface and anaerobes most prevalent at depth. (D) The [O_2_]/[H_2_S] ratio was strongly correlated with the inferred aerobe/anaerobe ratio on a *log-log* plot (Pearson *ρ* = 0.94, P < 10^−6^) such that estimating the redox gradient from sequencing data resulted in less than threefold error over ≈6 orders (mean multiplicative error of 2.6-fold).

To examine the quantitative relationship between local chemistry ([O_2_]) and physiology (O_2_ utilization), we next applied the AA 3-mer model along a well-studied natural O_2_ gradient (Fig. 2B). Due to its unique hydrography and intense density gradient, the surface of the Black Sea mixes poorly with deeper waters, leading to a sharp transition from oxia near the surface to anoxic and sulfidic habitats at depth (16, 17). As expected from these chemical transitions, the AA 3-mer classifier predicted sympathetic traces of O_2_ utilization with depth, with aerobic MAGs dominating near the surface and anaerobes in deeper waters below the mixed layer (Fig. 2C). This correspondence was also quantitative, with the [O_2_]/[H_2_S] ratio correlating strongly with inferred aerobe/anaerobe ratio, highlighting that chemical gradients might be “sensed” by statistical analysis of DNA sequences (Fig. 2D).

## Discussion

As a step towards establishing DNA sequencing as a multiplexed chemical sensor for natural environments, we established a stringent pipeline for building classifiers of microbial O_2_ utilization that works across the tree of life. Using this pipeline, we compared sophisticated annotation-based classifiers to simpler ones eliding genome annotation entirely. To our surprise, simple approaches based entirely on sequence content (e.g. amino acid triplet counts) performed nearly as well as intensive, annotation-based methods (Fig. 1C).

There are two mutually compatible explanations for the success of these naive approaches. First, O_2_ utilization is correlated with phylogeny to a degree (4) and *K*-mer counts are a proxy for phylogenetic proximity (18, 19). Indeed, NT *K*-mers are widely used to cluster DNA sequences during metagenome binning (20). Consistent with phylogeny predicting O_2_ utilization, we achieved useful prediction accuracy with a classifier trained on machine-learned embeddings of ribosomal 16S sequences (Fig. S4). Further, it has been argued that the sequence content of the whole genome adapts to O_2_ (21). If this is correct, we do not yet understand the biochemical logic underpinning this adaptation. For example, we attempted to summarize genomic sequence content with chemical metrics including elemental content (22) and the carbon redox state (21) of nucleotides and amino acids. A classifier based on these features displayed only 44% validation accuracy (Fig 1C). Nonetheless, a practical advantage of *K*-mer representations is that they contain both chemical and phylogenetic information, indicating that *K*-mers may be useful in simplifying model-based predictions of other complex physiologies (9).

Our exploration of diverse metagenomes indicated that the physical and chemical conditions of natural environments affects their sequence content in a legible way (Fig. 2). Indeed, we observed a quantitative correspondence between local chemistry ([O_2_]/[H_2_S]) and the inferred aerobe/anaerobe ratio in the Black Sea (Fig. 2D), suggesting that the inverse problem — estimating the concentrations of [O_2_] and other key molecules from sequences — is tractable. Perhaps sequencing data could serve as a “multi-sensor” of the local concentrations of several nutrients (e.g., O_2_, phosphate, nitrate, and sulfate) that play key roles in microbial growth and biogeochemical cycling. It is currently very challenging to characterize and monitor environmental chemistry at scale, whether for precision agriculture or for Earth system models. Our results here suggest that sequencing-as-sensing is a scalable way forward.

## Acknowledgements

The authors thank Daniel Dar, Jagoda Jabłońska, Ranjani Murali, and Jared Leadbetter for valuable discussions. This research was supported in part by the National Science Foundation under Grant No. NSF PHY-1748958. A.I.F was supported by the Jane Coffin Childs Memorial Fund for Medical Research. J.E.G. was supported by the Gordon and Betty Moore Foundation as Physics of Living Systems Fellows through grant number GBMF4513. A.J. acknowledges support from the Howard Hughes Medical Institute as a Hanna Gray Fellow (Grant #GT16787) and from the National Institute of Health through the UCSD FIRST program. W.W.F. acknowledges support from the Resnick Sustainability Institute, the Caltech Center for Evolutionary Sciences, and NSF NNA grant 2127442. This research was also sponsored by the Army Research Office and was accomplished under the Cooperative Agreement Number W911NF-22-2-0210 to D.K.N.

## Contributions

A.I.F., J.E.G, A.J., W.W.F. and D.K.N. designed the research. J.E.G., A.I.F., E.M.L., and A.J. wrote code and performed analysis. A.I.F. and J.E.G. wrote the manuscript with approval of all authors.

## Materials and Methods

### Code and data availability

Code and data are available on the following github repository: https://github.com/jgoldford/aerobot. Training data can be downloaded from Google Cloud using the provided download_training_data() function.

### Training and validation data

The dataset of Madin et al. (8) was mapped onto the Genome Taxonomy Database (GTDB release r207 (23)) to produce a collection of genomes and metagenomes with known modes of dioxygen utilization. We generated three classes of labels for each genome using the following rules: we labeled annotations “Anaerobe”, “Obligate anaerobe” as “Anaerobe”, “Facultative”, “Facultative anaerobe” as “Facultative”, and “Aerobe”, “Microaerophilic”, “Obligate aerobe” as “Aerobe”. Validation data was drawn from (2) where a curated set of phylogenetically-balanced bacteria and archaea was used to investigate the relationship between O_2_ utilizing enzymes and dioxygen requirement. Validation set genomes and coding sequences were retrieved using RefSeq IDs provided in (2). Genomes were processed with a custom Python pipeline to extract features (e.g. nucleotide tetramers). Genome annotation was performed using kofamscan (14) and protein embedding was performed with the protein language model ProtT5-XL-uniref50 (12). Genomes were represented as averages of protein coding sequence embeddings for the “genome embedding” model in Fig. 1C.

### Model training and validation

We developed a common pipeline to evaluate the predictive power of 17 feature sets (Fig. S2). Annotation-free feature sets included the number of predicted open reading frames (“gene count”), counts of genomic DNA *K*-mers (lengths 1-5), counts of coding sequence amino acid *K*-mers (lengths 1-3), a manually selected list of simple chemical features of nucleotide and amino acid sequences in each genome (“chemical features”), and genome embeddings. Chemical features included the number of open reading frames, the genomic GC content, the number of carbon, nitrogen, oxygen and sulfur atoms (22) in protein and RNA coding sequences as well as the average redox state of carbon (*Z*_*C*_) in those same sequences (21). The “gene embedding” approach requires no annotation but adds a computationally intensive step as it involves a forward pass through a large language model (12). Annotation-full feature sets included per-genome counts of KEGG orthogroups (“All gene families”), counts of O_2_ utilizing enzymes, a curated set of the 5 most-predictive O_2_-utilizing enzyme activities identified in (2), counts of O_2_-utilizing terminal oxidases (e.g. heme-copper O_2_ reductases), and two 1D feature sets: the number of O_2_ utilizing genes and the fraction of genes that are O_2_ utilizing. The 5 most predictive O2 utilizing enzymes were coproporphyrinogen III oxidase (enzyme commission number EC 1.3.3.3), nitric-oxide synthase (EC 1.14.14.47), stearoyl-CoA desaturase (EC 1.14.19.1), aklavinone 12-hydroxylase (EC:1.14.13.180), and FAD-dependent urate hydroxylase (EC 1.14.13.113). We then applied *L*_2_-regularized logistic regression (Python *sklearn* package; regularization strength set to *C*=100) to estimate the mapping between features and labels. We then calculated balanced accuracy across the training and validation sets (*sklearn*.*metrics* package).

To predict O_2_ utilization from 16S rRNA DNA sequence, we fine-tuned the DNA Language Model GenSLM (24). For genomes with NCBI accession for 16S rRNA genes, we extracted the V34 region and embedded this into a 512-dimensional vector using GenSLM. NCBI accessions were not available for all genomes in (8), meaning that 16S sequences could not be ascertained in all cases. This resulted in a dataset of *n* = 899 variable regions from genomes with known oxygen requirements. We randomly partitioned the data into training sequences (*n* = 720), validation sequences (*n* = 80), and testing sequences (*n* = 99). We constructed a classification layer on top of GenSLM in PyTorch, varying only the weights in this additional layer during training. The model was trained with a learning rate of 0.01, a batch size of 32, and for 200 epochs using the Adam optimizer. The final model was chosen via early stopping at epoch 151, which corresponded to the model with the highest balanced accuracy for the validation set during training. Note that this model uses different training and test sets than models trained on full genomes, making direct comparison challenging. As such, these model results are not directly comparable with those in Figs. 1 and S2, so they are presented separately in Fig. S4.

### Black sea analysis

The relative abundances of metagenome-assembled genomes (MAGs) from metagenomic samples in the Black Sea (16) were generated in prior studies (25). Briefly, metagenomic samples were aligned to the previously assembled MAGs using *bbmap*. Alignments with a mapping quality above 10 were retained, converted to BAM format, sorted, and indexed using *samtools*. The relative abundance of each MAG was determined by the fraction of reads mapped to it, as summarized by samtools *idxstats*. This process was automated via a Python script, utilizing samtools v1.8 and bbmap.sh, and executed on the Resnick High-Performance Computing Center cluster at Caltech.

### Earth Microbiome Project (EMP) analysis

O_2_ utilization phenotypes of EMP metagenome assembled genomes (MAGs) from (15) were classified using the AA 3-mer model. Since EMP projects are categorized with an ad hoc nomenclature describing the environment sampled, we manually mapped tags to a simplified set of categories. For analysis, we removed MAGs with less than 50% estimated completeness, considered only samples from which at least 10 MAGs were assembled and only environmental labels (e.g. “rhizosphere”) for which at least 10 samples were available. This left 1598 samples and 31,279 MAGs under consideration. The data presented in Fig. 2C gives the fraction of MAGs that are inferred to be aerobes, anaerobes, and facultative in each habitat for samples meeting these criteria.

## Supplementary Figures

**Fig. S1:**
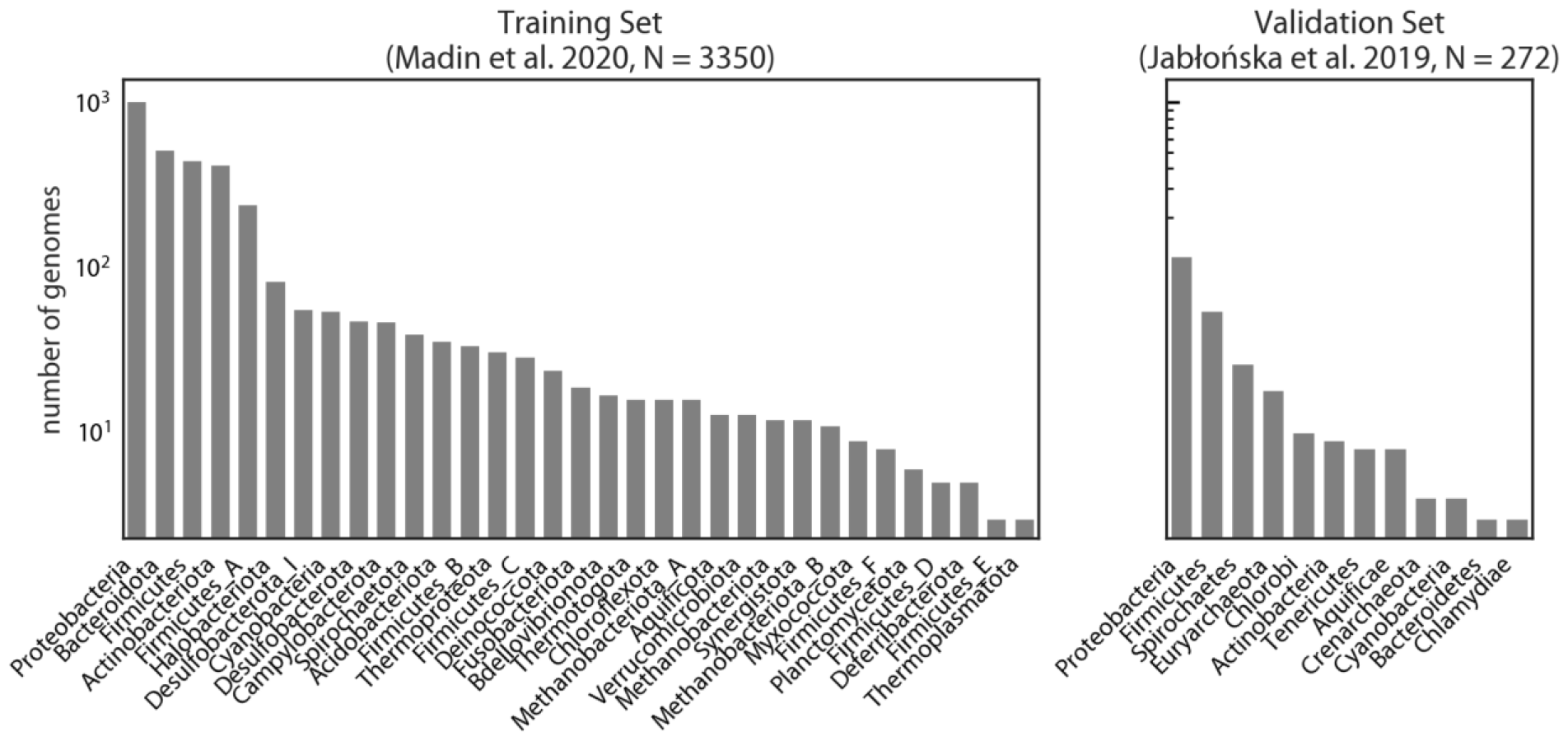
The phylogenetic coverage of the training and validation genome sets expressed at the phylum level. Panels display counts from phyla with at least 3 representatives in the training (A) and validation (B) dataset.

**Fig. S2:**
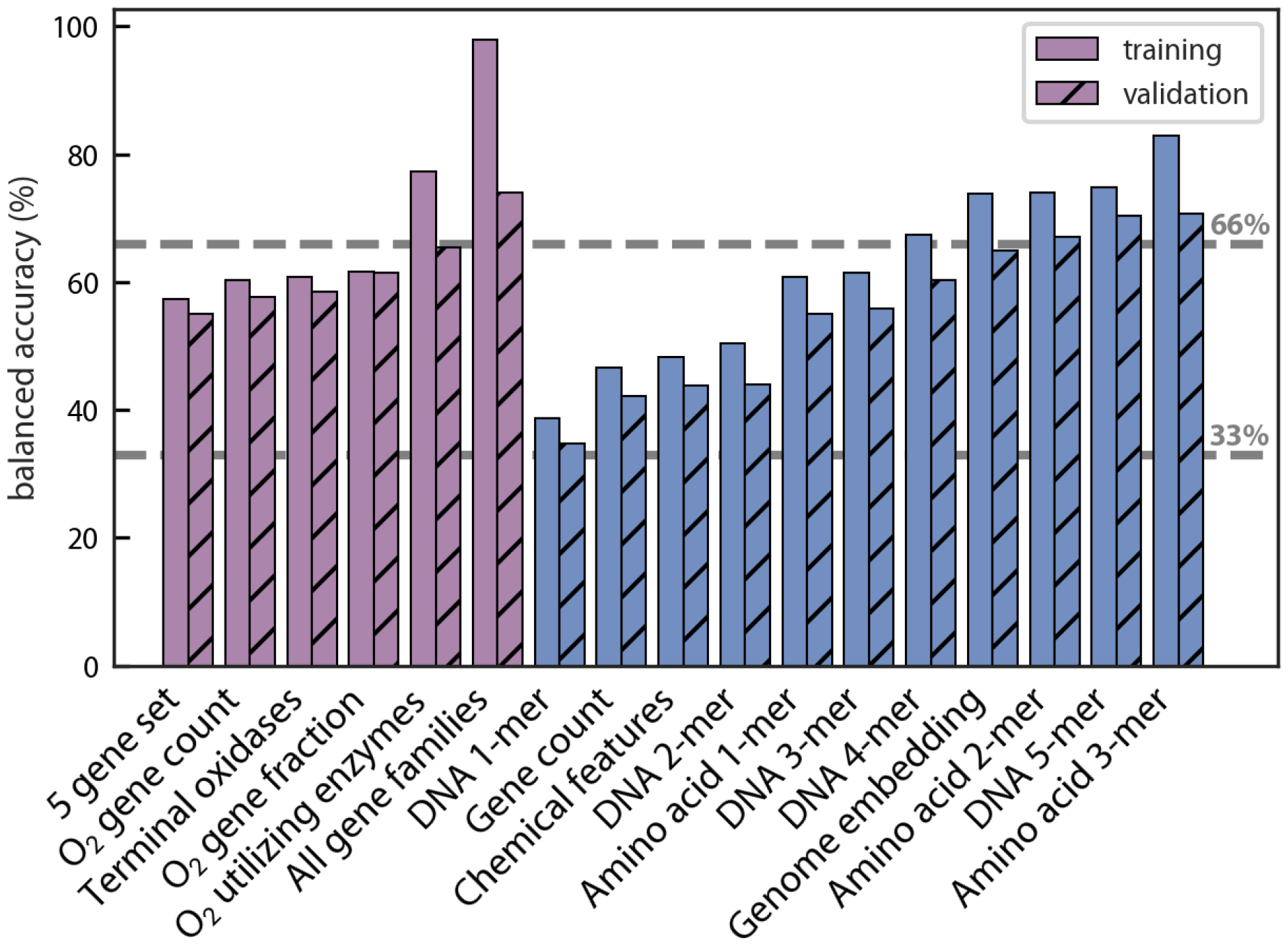
Training and test-balanced accuracies for all models trained. Models are grouped by whether or not they are annotation-full (purple, left) or annotation-free (blue, right). Within groups, they are ordered left-to-right by increasing balanced accuracy on the validation set (hatched bars). Notice that three of the top five performing approaches were annotation-free: genomic DNA 5-mers, amino acid 3-mers, and genome embeddings. The dashed gray lines in the background mark the random guessing threshold (33% balanced accuracy) and twice that threshold (66% balanced accuracy) for ternary classification. See Methods for detailed description of the individual models.

**Fig. S3:**
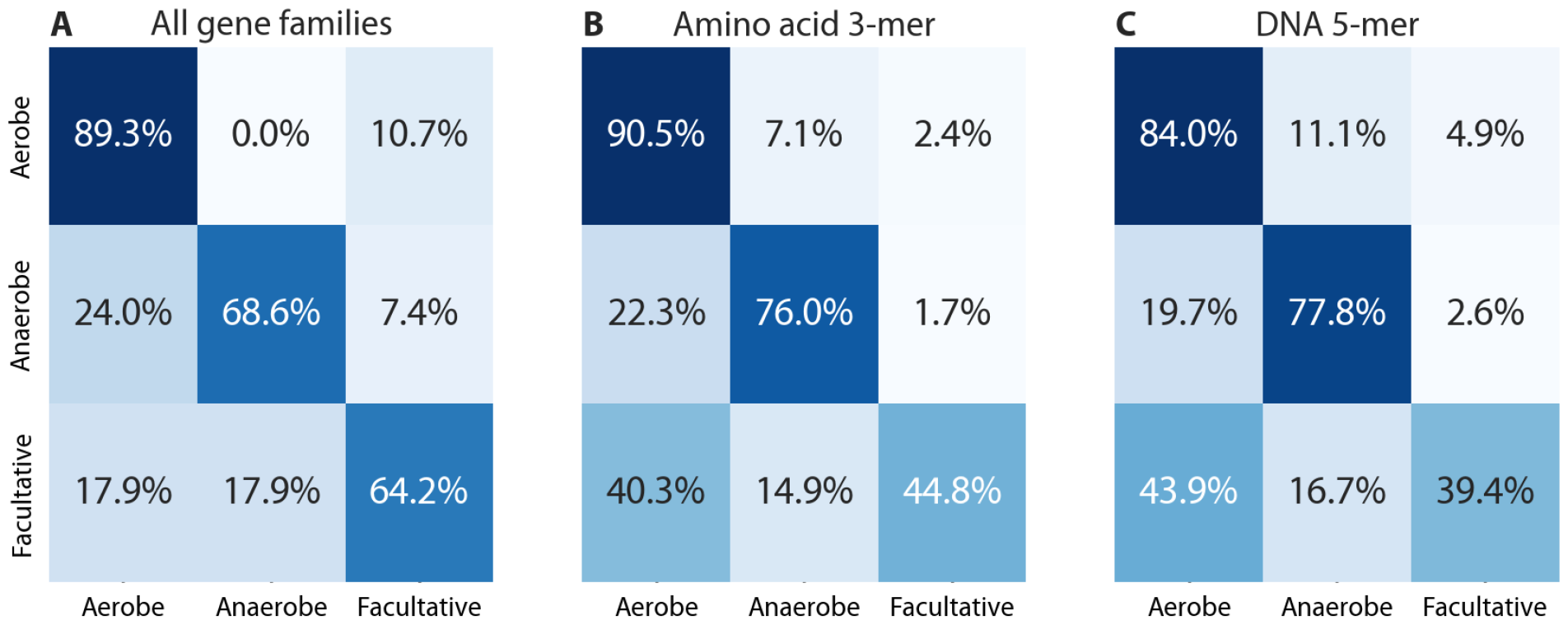
Confusion matrices for three top-performing models. Confusion matrices depict the per-class accuracies. Notice that, in all cases, facultative organisms were the most difficult to accurately classify. (A) All gene families model; 74% total balanced validation accuracy, (B) AA 3-mer model; 70%, and (C) DNA 5-mer model (67%) as described in Methods.

**Fig. S4:**
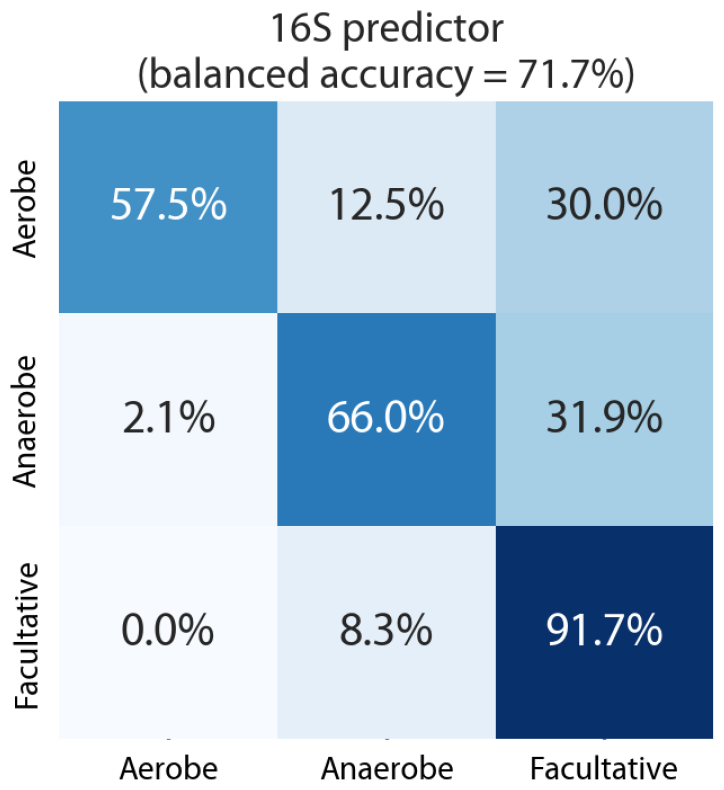
A predictor based on embedding 16S sequences performs similarly to other approaches, but with a distinct error profile. We used machine-learning driven DNA sequence embedding to develop a classifier based on 16S rRNA sequences. Due to constraints associated with our validation set, however, balanced accuracy is here calculated over a smaller, randomized testing set containing 99 examples (Methods). Accuracy values are therefore not directly comparable to Figs. 1, S2 or S3.

**Fig. S5:**
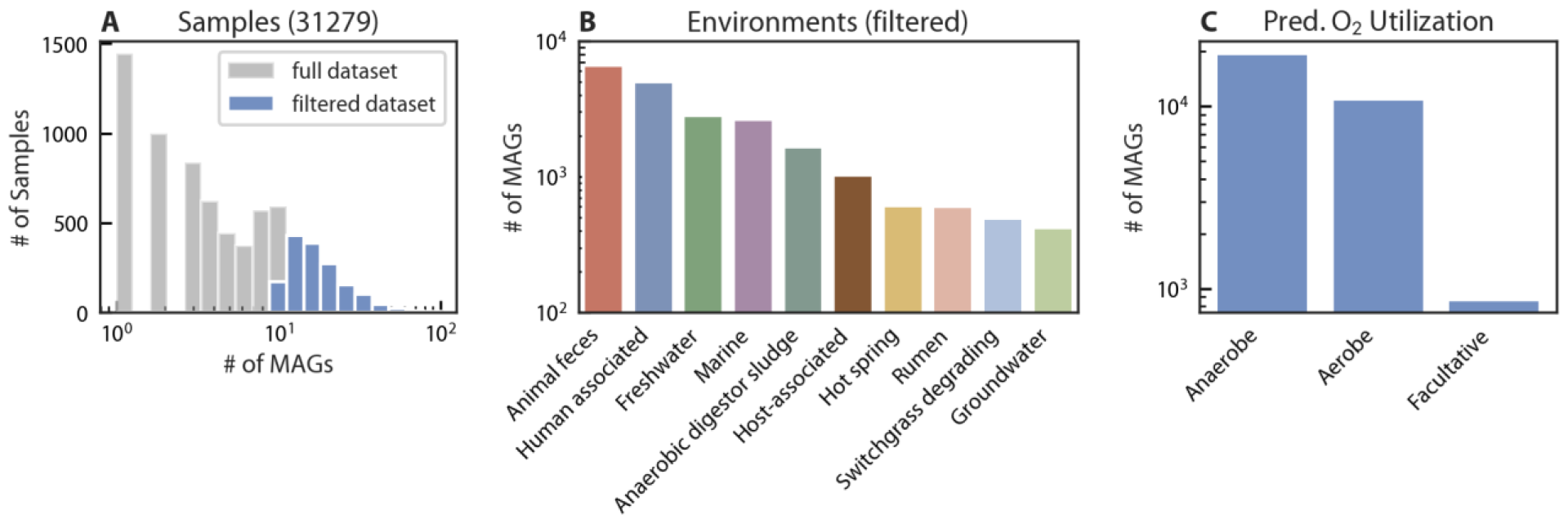
Summary of the Earth Microbiome Project metagenomic compendium. (A) The number of metagenome-assembled genomes (MAGs) associated with individual samples was non-uniform. Samples with fewer than 10 associated MAGs were not considered in subsequent analyses. (B) Sample environments were broadly categorized so that samples could be grouped and compared by environment. Here we show the environments with the largest number of associated samples. (C) Counts of O_2_ utilization as inferred by running the AA 3-mer model on Earth Microbiome Project MAGs.

## References

1. Fischer WW, Valentine JS. 2019. How did life come to tolerate and thrive in an oxygenated world? Free Radical Biology and Medicine.

2. Jabłońska J, Tawfik DS. 2019. The number and type of oxygen-utilizing enzymes indicates aerobic vs. anaerobic phenotype. Free Radic Biol Med 10.1016/j.freeradbiomed.2019.03.031.

3. Lu Z, Imlay JA. 2021. When anaerobes encounter oxygen: mechanisms of oxygen toxicity, tolerance and defence. Nat Rev Microbiol 10.1038/s41579-021-00583-y.

4. Nielsen DA, Fierer N, Geoghegan JL, Gillings MR, Gumerov V, Madin JS, Moore L, Paulsen IT, Reddy TBK, Tetu SG, Westoby M. 2021. Aerobic bacteria and archaea tend to have larger and more versatile genomes. Oikos 10.1111/oik.07912.

5. Hackmann TJ, Zhang B. 2023. The phenotype and genotype of fermentative prokaryotes. Sci Adv 9:eadg8687.

6. Weber Zendrera A, Sokolovska N, Soula HA. 2021. Functional prediction of environmental variables using metabolic networks. Sci Rep 11:12192.

7. Edirisinghe JN, Goyal S, Brace A, Colasanti R, Gu T, Sadhkin B, Zhang Q, Kamimura R, Henry CS. 2023. Machine Learning-Driven Phenotype Predictions based on Genome Annotations. bioRxiv.

8. Madin JS, Nielsen DA, Brbic M, Corkrey R, Danko D, Edwards K, Engqvist MKM, Fierer N, Geoghegan JL, Gillings M, Kyrpides NC, Litchman E, Mason CE, Moore L, Nielsen SL, Paulsen IT, Price ND, Reddy TBK, Richards MA, Rocha EPC, Schmidt TM, Shaaban H, Shukla M, Supek F, Tetu SG, Vieira-Silva S, Wattam AR, Westfall DA, Westoby M. 2020. A synthesis of bacterial and archaeal phenotypic trait data. Sci Data 7:170.

9. Gralka M, Pollak S, Cordero OX. 2023. Genome content predicts the carbon catabolic preferences of heterotrophic bacteria. Nat Microbiol 10.1038/s41564-023-01458-z.

10. Kislyuk A, Bhatnagar S, Dushoff J, Weitz JS. 2009. Unsupervised statistical clustering of environmental shotgun sequences. BMC Bioinformatics 10:316.

11. Bray NL, Pimentel H, Melsted P, Pachter L. 2016. Near-optimal probabilistic RNA-seq quantification. Nat Biotechnol 34:525–527.

12. Elnaggar A, Heinzinger M, Dallago C, Rehawi G, Wang Y, Jones L, Gibbs T, Feher T, Angerer C, Steinegger M, Bhowmik D, Rost B. 2022. ProtTrans: Toward Understanding the Language of Life Through Self-Supervised Learning. IEEE Trans Pattern Anal Mach Intell 44:7112–7127.

13. Lin Z, Akin H, Rao R, Hie B, Zhu Z, Lu W, Smetanin N, Verkuil R, Kabeli O, Shmueli Y, Dos Santos Costa A, Fazel-Zarandi M, Sercu T, Candido S, Rives A. 2023. Evolutionary-scale prediction of atomic-level protein structure with a language model. Science 379:1123–1130.

14. Aramaki T, Blanc-Mathieu R, Endo H, Ohkubo K, Kanehisa M, Goto S, Ogata H. 2020. KofamKOALA: KEGG Ortholog assignment based on profile HMM and adaptive score threshold. Bioinformatics 36:2251–2252.

15. Nayfach S, Roux S, Seshadri R, Udwary D, Varghese N, Schulz F, Wu D, Paez-Espino D, Chen I-M, Huntemann M, Palaniappan K, Ladau J, Mukherjee S, Reddy TBK, Nielsen T, Kirton E, Faria JP, Edirisinghe JN, Henry CS, Jungbluth SP, Chivian D, Dehal P, Wood-Charlson EM, Arkin AP, Tringe SG, Visel A, IMG/M Data Consortium, Woyke T, Mouncey NJ, Ivanova NN, Kyrpides NC, Eloe-Fadrosh EA. 2021. A genomic catalog of Earth’s microbiomes. Nat Biotechnol 39:499–509.

16. Villanueva L, von Meijenfeldt FAB, Westbye AB, Yadav S, Hopmans EC, Dutilh BE, Damsté JSS. 2021. Bridging the membrane lipid divide: bacteria of the FCB group superphylum have the potential to synthesize archaeal ether lipids. ISME J 15:168–182.

17. Özsoy E, Ünlüata Ü. 1997. Oceanography of the Black Sea: A review of some recent results. Earth-Sci Rev 42:231–272.

18. Bernard G, Ragan MA, Chan CX. 2016. Recapitulating phylogenies using k-mers: from trees to networks. F1000Res 5:2789.

19. Bussi Y, Kapon R, Reich Z. 2021. Large-scale k-mer-based analysis of the informational properties of genomes, comparative genomics and taxonomy. PLoS One 16:e0258693.

20. Dick GJ, Andersson AF, Baker BJ, Simmons SL, Thomas BC, Yelton AP, Banfield JF. 2009. Community-wide analysis of microbial genome sequence signatures. Genome Biol 10:R85.

21. Dick JM, Meng D. 2023. Community- and genome-based evidence for a shaping influence of redox potential on bacterial protein evolution. mSystems e0001423.

22. Shenhav L, Zeevi D. 2020. Resource conservation manifests in the genetic code. Science 370:683–687.

23. Parks DH, Chuvochina M, Waite DW, Rinke C, Skarshewski A, Chaumeil P-A, Hugenholtz P. 2018. A standardized bacterial taxonomy based on genome phylogeny substantially revises the tree of life. Nat Biotechnol 36:996–1004.

24. Zvyagin M, Brace A, Hippe K, Deng Y, Zhang B, Bohorquez CO, Clyde A, Kale B, Perez-Rivera D, Ma H, Mann CM, Irvin M, Gregory Pauloski J, Ward L, Hayot V, Emani M, Foreman S, Xie Z, Lin D, Shukla M, Nie W, Romero J, Dallago C, Vahdat A, Xiao C, Gibbs T, Foster I, Davis JJ, Papka ME, Brettin T, Stevens R, Anandkumar A, Vishwanath V, Ramanathan A. 2022. GenSLMs: Genome-scale language models reveal SARS-CoV-2 evolutionary dynamics. bioRxiv.

25. Goldford JE, Murali R, Valentine JS, Fischer WW. 2023. Metabolic evolution of pyranopterindependent biochemistry. bioRxiv.

